# A causal role for mouse superior colliculus in visual perceptual decision-making

**DOI:** 10.1101/835066

**Authors:** Lupeng Wang, Kerry McAlonan, Sheridan Goldstein, Charles R. Gerfen, Richard J. Krauzlis

## Abstract

The superior colliculus (SC) is arguably the most important visual structure in the mouse brain and is well-known for its involvement in innate responses to visual threats and prey items. In other species, the SC plays a central role in voluntary as well as innate visual functions, including crucial contributions to selective attention and perceptual decision-making. In the mouse, the possible role of the SC in voluntary visual choice behaviors has not been established. Here, we demonstrate that the mouse SC plays a causal role in visual perceptual decision-making by transiently inhibiting SC activity during an orientation-change detection task. First, unilateral SC inhibition induced spatially specific deficits in detection. Hit rates were reduced and reaction times increased for orientation changes in the contralateral but not ipsilateral visual field. Second, the deficits caused by SC inhibition were specific to a temporal epoch coincident with early visual burst responses in the SC. Inhibiting SC during this 100-ms period caused a contralateral detection deficit, whereas inhibition immediately before or after did not. Third, SC inhibition reduced visual detection sensitivity. Psychometric analysis revealed that inhibiting SC visual activity significantly increased detection thresholds for contralateral orientation changes. In addition, effects on detection thresholds and lapse rates caused by SC inhibition were larger in the presence of a competing visual stimulus, indicating a role for the mouse SC in visual target selection. Together, our results demonstrate that the mouse SC plays a crucial role in voluntary visual choice behaviors.

**Significance statement:** The mouse superior colliculus has become a popular model for studying the circuit organization and development of the visual system. Although the SC is a fundamental component of the visual pathways in mice, its role in visual perceptual decision-making is not clear. By investigating how temporally precise SC inhibition influenced behavioral performance during a visually guided orientation change detection task, we identified a 100-ms temporal epoch of SC visual activity that is crucial for the ability of mice to detect behaviorally relevant visual changes. In addition, we found that SC inhibition also caused deficits in visual target selection. Thus, our findings highlight the importance of the SC for visual perceptual choice behavior in the mouse.

## Introduction

The mouse has emerged as a powerful animal model for investigating visual neural circuits, thanks to the abundant genetic tools available in this species (Huberman and Niell, 2011). We might now have more knowledge about the early visual system in mice at molecular, neuronal and circuit levels than for any other sensory system in any species (Seabrook et al., 2017), but how mice use visual information to guide voluntary actions has been less studied. Among all visual structures in mice, the superior colliculus (SC) is one of the most important. The mouse SC gets direct inputs from almost all retinal ganglia cells (Ellis et al., 2016) and is also a major output target of the visual cortex (Cang and Feldheim, 2013). Thus, understanding the functional role of the SC is crucial for understanding vision in mice.

Early visual processing in the SC of mice is important for fast innate visually guided behaviors, such as avoiding predators (Shang et al., 2015; Wei et al., 2015) and approaching prey (Hoy et al., 2016; Shang et al., 2019). Neurons in the medial SC of the mouse detect looming visual threats overhead and trigger avoidance behaviors, including escape responses mediated through projections to the dorsal periaqueductal gray (Evans et al., 2018) and freezing responses mediated through the basolateral amygdala (Shang et al., 2015; Wei et al., 2015). Lateral SC neurons in mice detect prey items in the lower visual field and illicit hunting and approach responses through projections to zona incerta (Shang et al., 2019). Conserved across all species, the SC also sends extensive projections to brainstem and spinal cord motor neurons that generate visually guided orienting responses (Dean et al., 1988; Gandhi and Katnani, 2011; Wurtz and Albano, 1980).

In several other species, the SC also plays a crucial role in voluntary, flexible and learned visual perceptual functions, such as perceptual decision-making (Boehnke and Munoz, 2008; Herman et al., 2018), visual target selection (Carello and Krauzlis, 2004; Horvitz and Newsome, 1999) and spatial attention (Krauzlis et al., 2013). To investigate the visual perceptual functions of the SC, it is especially important to dissociate SC’s role in sensory processing from its role in movement preparation. Lesions or reversible inactivation of the SC can cause behavioral impairments for visual stimuli presented in the affected visual field, such as deficits in saccade generation in primates, seed-collecting in hamsters (Schneider, 1969) and nose-poking in rats (Dean and Redgrave, 1984). The spatially specific deficits observed in these studies could be due to motor impairments in the ability to orient toward or away from the affected hemifield, rather than impairments in perceptual processing. This ambiguity can be resolved by using a response that does not involve spatial orienting, for example, by having a monkey press a button or release a joystick (Herman et al., 2018; Lovejoy and Krauzlis, 2017). Another possibility is to use optogenetic methods to limit the suppression of SC activity to specific sensory or motor preparatory epochs of the task, but these methods are not yet reliable in monkeys, and mice have not yet been tested in appropriate tasks.

Here we investigated the role of mouse SC in perceptual decision-making by training mice to lick a center spout to report the detection of a visual orientation change and used optogenetics to inhibit neuronal activity in the SC during specific temporal windows. We found that unilateral inhibition of SC activity decreased accuracy and increased reaction time for detecting near-threshold orientation changes in the contralateral hemifield but left performance in the ipsilateral hemifield unaffected, consistent with the visual representation in the SC. These spatially specific deficits were only found when SC activity was inhibited during a 100-ms interval that coincided with transient visual response in the SC. Together, our results demonstrate that visual activity in the mouse SC is crucial for detecting behaviorally relevant events during perceptual decisions.

## Material and Methods

### Animals

All procedures were conducted on vgat-ires-cre (JAX stock # 028862, Jackson Laboratory, Bar Harbor, ME) and wild-type C57BL/6J mice (JAX stock # 000664). Vgat-ires-cre mice were derived from homozygotes mated with wild-type C57BL/6J mice, producing heterozygotes littermates. All the transgenic animals used in the study were heterozygotes. The mice were housed in a 12:12 reversed day-night cycle, with lights off at 9 am, and all experimental procedures and behavioral training were done in the lights-off portion of the cycle (9am-9pm). Male and female mice weighing 18-25 grams were surgically implanted at age 6-8 weeks and then used in experiments for up to ∼9 months. All the mice were in group housing (2-4 cage mates) prior to the surgical procedure, and subsequently singly housed after the implant surgery. All experimental procedures and animal husbandry were approved by the NIH Institutional Animal Care and Use Committee (IACUC) and complied with Public Health Service policy on the humane care and use of laboratory animals.

### Viral vectors

Double-floxed inverted (DIO) recombinant Adeno-associated viral vectors (AAV2) was used to express light-gated channel channelrhodopsin (hChR2(H134R)-eYFP) in Cre-expressing GABAergic neurons in the superior colliculus. The double-floxed reverse hChR2-eYFP cassette was driven with an EF-1a promoter and WPRE to enhance expression. The recombinant AAVs vector was packaged by the University of North Carolina viral core (titer of 4×10^12^ particles/ml).

### Stereotaxic surgery

Each mouse was injected with virus unilaterally and implanted with a head-holder and optic fiber during a single surgical procedure. During the surgery, animals were anesthetized with isoflurane (4% induction, 0.8-1.5% maintenance) and secured by a stereotaxic frame with ear bars (Kopf Instruments). Dexamethasone (1.6 mg/kg) was administered to reduce inflammation. A feedback-controlled heating pad (FHC) was used to maintain the body temperature at 37°C, and artificial tears were applied to the eyes to prevent them from drying. After the animal’s head was leveled in the stereotaxic frame, a scalp incision was made along the midline, followed by a small craniotomy for virus injection and optic fiber implantation. The coordinates for the unilateral virus injection were ±0.8∼1.1mm from midline (M-L axis), −3.65∼-4.0mm from Bregma (A-P axis) and 1.8-2.1mm ventral (D-V axis), based on a standard mouse brain atlas {Paxinos:2004ts}. Each mouse was injected with 0.1-0.2 microliter of virus at a flow rate of 50 nl/min, using a manual microinjector (Sutter Instrument) with 30μm-tip pulled glass pipettes. An optic fiber (200μm core) together with its ceramic ferrule base (Plexon Inc.) were subsequently inserted at the injection coordinates with the fiber tip located at 0.3∼0.5mm above the injection center. For the mouse used in recording SC visual responses, a custom 16-wire microwire bundle with Microdrive (Innovative Neurophysiology, Durham, NC) was implanted over the cranial window, and lowered into the SC. A custom-designed titanium head post for head-fixing was positioned and secured to the skull together with the ferrule using Metabond (Parkell Inc.). The skin wound edge was then closed with sutures or tissue adhesive (3M Vetbond). After surgery, mice received subcutaneous ketoprofen (1.85mg/kg) daily for up to three days to ease discomfort. Fiber placement and viral expression were validated histologically in all mice after completion of data collection.

### Food control

After mice recovered from surgery and returned to above 95% of their pre-surgery weight (typically within 7-9 days), they were placed on a food control schedule. Mice had free access to water, but their intake of dry food was controlled, and they were allowed to augment their dietary intake by access to a nutritionally complete 8% soy-based infant formula (Similac, Abbott, IL). Overall food intake was regulated to maintain at least 85% of their free-feeding body weight, and the health status of each mice was monitored daily throughout the study. Mice were initially acclimatized to handling procedures by having their heads gently restrained while receiving the soy-based fluid under manual control via a sipper tube. After the initial exposure to soy-based fluid, we more securely head-fixed the animal and continued manual delivery. Once mice were adapted to these procedures, we switched to automatic delivery of fluid under computer control in the behavioral apparatus.

### Behavioral apparatus

The behavioral apparatus consisted of a custom-built booth that displayed visual stimuli to the mouse coupled to their locomotion. The mouse was head-fixed in the center of the apparatus, positioned atop a polystyrene foam wheel (20-cm diameter) that allowed natural walking or running movements along a linear path. An optical encoder (Kübler) was used to measure the rotation of the wheel. The front walls of the booth incorporated a pair of LCD displays (ViewSonic VG2439) positioned at 45° angles from the animal’s midline such that each display was centered on either the right or left eye and subtended ∼90° horizontal by ∼55° vertical of the visual hemifield, with a viewing distance of 27.5 cm. The interior of the booth was lined with sound absorbent material to reduce acoustic noise. The experiments were controlled by a computer using a modified version of the PLDAPS system (Eastman and Huk 2012). Our system omitted the Plexon device, but included a Datapixx peripheral (Vpixx Technologies, Saint-Bruno, QC, Canada) and the Psychophysics Toolbox extensions (Brainard, 1997; Pelli, 1997) for Matlab (The Mathworks, Natick, MA, USA), controlled by Matlab-based routines run on a Mac Pro (Apple, Cupertino, CA, USA). The Datapixx device provided autonomously timed control over analog and digital inputs and outputs and synchronized the display of visual stimuli generated using the Psychophysics Toolbox.

A reward delivery spout was positioned near the snout of the mouse; lick contacts with the spout were detected by a piezo sensor (Mide Technology Co., Medford, MA, USA) and custom electronics. Each reward was a small volume (5-10 μl) of an 8% solution of soy-based infant formula (Similac, Abbott, IL) delivered by a peristaltic pump (Harvard Apparatus) under computer and Datapixx control. Airpuff aversive stimuli were delivered through a second spout located slightly above the reward spout, and controlled through solenoids (Parker Hannifin, Cleveland, OH, USA). The temperature inside the apparatus maintained between 70-80° F.

### Visual detection tasks

The detection task was modified from our previously published study (Wang et al., 2018). Animals were run in experiments on consecutive days and each session produced 300-900 trials. Experiments were organized in blocks of randomly shuffled, interleaved trials, and each trial consisted of a sequence of epochs that the mouse passed through by walking or running on the wheel. Each epoch was defined by the particular stimuli presented on the visual displays, and the duration of each epoch was determined by the time that it took for the mouse to travel a randomized distance on the wheel, typically several seconds based on the running speed of the mice (median: 85.6 cm/s).

In all of the experiments, each trial followed a standard sequence of four epochs. The average luminance across each visual display in all epochs was 4-8 cd/m^2^. In the first epoch (“noise”), the uniform gray of the inter-trial interval was changed to pink visual noise with an RMS contrast of 3.3%; this epoch was presented for a distance of 0.9-1.8 cm (i.e., 0.007-0.03 s). In the second epoch (“cue”), a contrast annulus was added to the pink noise, center in either left or right visual display. The annulus consisted of a sinusoidal concentric function, with radius and width matched to the Gabor patch subsequently shown. This second epoch lasted for 46-92 cm (0.36-1.55 s). In the third epoch (“delay”), it depended on the trial type. If it was a single-patch trial, a vertically oriented Gabor patch was added to the pink noise, centered in either the left or right visual display. The Gabor patch consisted of a sinusoidal grating (95% Michelson contrast) with a spatial frequency of 0.1 cycles per degree, a value chosen based on the visual spatial acuity of mice (Sinex et al., 1979), modulated by a Gaussian envelope with full width at half-maximum of 18° (σ = 7.5°). The phase of the grating was not fixed, but throughout the trial was incremented in proportion to the wheel rotation with every monitor refresh, so that the sinusoidal pattern was displaced on the screen by approximately the same distance that the mouse traveled on the wheel; the Gabor patch on the left (right) drifted leftward (rightward), consistent with optic flow during locomotion. If it was a double-patch trial, two Gabor patches with the same properties as in single-patch trials appeared on the opposite sides of the visual displays. This third epoch lasted for 107-214 cm (0.84-3.6 s). The visual stimuli in the fourth epoch depended on whether the trial was a “change” or “no change” condition; the two types were equally likely and randomly interleaved within a block, and both lasted 77-154 cm (0.61-2.6 s). On single-patch change trials, the Gabor patch changed its orientation at the onset of the fourth epoch. On double-patch change trials, the Gabor patch on the same side as the cue ring changed its orientation at the onset of the fourth epoch. The direction of the orientation change observed mirror symmetry: if the left (right) Gabor changed, it rotated clockwise (counter-clockwise). On no-change trials, neither Gabor patch changed its orientation, so that the fourth epoch unfolded as a seamless extension of the previous two-patch epoch.

The task of the mouse was to lick the spout when he or she detected a change in the orientation of the Gabor patch and to otherwise withhold from licking. Mice were required to lick within a 500-ms response window starting 300 ms after the orientation change in order to score a “hit” and receive a fluid reward. The initial 300 ms no-lick period was applied to most mice, except for a few mice that had faster reaction times, for whom the response windows were shifted earlier by 50 ms. If the mouse failed to lick within this window after an orientation change, the trial was scored as a “miss” and no reward was given but no other penalty was applied. On no-change trials, if the mouse licked within the same response window aligned on the transition to epoch 4, the trial was scored as a “false alarm”; if they correctly withheld from licking, the trial was scored as a “correct reject”. At the end of correct reject trials, the trial was extended to include an additional “safety-net epoch” in which the initially appearing Gabor underwent a supra-threshold (30°) orientation change and the mouse could receive a reward by licking within a comparable response window. The point of this safety-net epoch was to maintain motivation by rewarding the mouse for correct behavior without violating the task rule that they should lick only for orientation changes. False alarms and premature licks before the response window led to timeouts and possible air-puff penalties; well-trained mice usually committed fewer than 10% of such errors.

All trials followed the standard 4-epoch trial sequence and were organized into alternating blocks in which the orientation change occurred in either the left or right Gabor patch. We counterbalanced the frequency of trials with and without orientation changes, and with and without stimulation, in order to minimize possible behavioral biases related to frequency matching. Each block of trials with cue on the same side included two sub-blocks, started with 60 single-patch trials followed with 60 double-patch trials. Blocks of left and right cue trials were randomly interleaved within each experimental session.

For experiments with 300-ms SC inhibition, we used a single value of 15° for the orientation change. For experiments with orange LED stimulation, we used a 12° orientation change. Each of the 4 possible combinations of two factors (change/no-change, stimulation/no stimulation) comprised 25% of the total trials in each block.

For experiments with 3 epochs of SC inhibition, we used a single value of 12° for the orientation change. Each sub-block of 60 trials with cue on the same side contained 25% trials with orientation change and no inhibition, 25% trials without orientation change and no inhibition, 25% trials with orientation change and SC inhibition drawn equally from 3 epochs, and 25% trials without orientation change and the same 3 epochs inhibition. All different trial types within the same sub-block were randomly interleaved.

For the psychometric curve experiments, each sub-block of 60 trials contained 25% with an orientation change drawn equally from five possible values (4, 7, 11, 15, 20°) and with no inhibition, 25% with the same five orientation changes with SC inhibition during epoch 2, 25% with no change and no inhibition, 25% with no change and inhibition during epoch 2. Because only some of the orientation changes could be reliably detected, a safety-net epoch was added to the end of miss trials, but with a long duration to discourage using it as a default response. We recorded 8-10 sessions for each mouse to obtain at least 60 repetitions for each of the five orientation changes. The experimenters were blind to the side of SC inhibition of animals, and the order of presentation for the different trial types in each experiment was also randomized.

### Electrophysiological recording

Visual responses of SC neurons were recorded in the male C57BL/6J mouse implanted with a moveable 16-wire microwire bundle (Innovative Neurophysiology, NC). Electrophysiological signals were acquired through an RZ5D processor (Tucker-Davis Technologies) with spikes filtered between 0.3 to 7 kHz and sampled of 25k Hz. Single units were sorted offline using KiloSort (Pachitariu et al., 2016). The SC surface was determined as the first visual responses encountered while advancing the microwire bundle. All single unit data were collected from depth within 2mm from the estimated SC surface.

### Monitoring eye movements

A high speed, 240 Hz CCD camera (ISCAN, Inc) was used to monitor the eye position of head-fixed mice during the 3-epoch SC inhibition experiments. We imaged an area of 1.5 mm x 3 mm with a macro lens (ISCAN, Inc) centered on the eye. Four infrared light-emitting-diodes (LEDs, 940nm) were used to illuminate the eye. Commercially available acquisition software (ETL-200, ISCAN) was used to determine the center and boundary of the pupil. Eye position was obtained by subtracting the center of corneal reflection from the pupil center to compensate any translational movement of the eye at the imaging plane. The pupil displacement in 2-D image was converted to a rotation angle based on estimated eyeball radius (1.25mm) from model C57bl/6 mice (Sakatani and Isa, 2004; Stahl, 2004). Eye velocity was then obtained by applying a low-pass differentiating filter to eye position traces (−3db rolloff at 54 Hz). Saccade detection was done in custom graphic user interface written in MatLab script, following a previously described algorithm (Krauzlis and Miles 1996), thresholded at minimum velocity of 70°/s, minimum acceleration of 3000°/s^2^, and minimum duration of 15-ms, and all eye traces were manually inspected to remove blink artifacts. Saccade probability was calculated as the fraction of trials within each time bin that were marked as containing a saccade.

### Histology

Mice were euthanized with CO_2_ (1.0 LPM), after which they were transcardially perfused with ice-cold saline followed by phosphate buffered saline with 4% paraformaldehyde (PFA). Their brains were removed and stored in the 4% PFA solution overnight, and then transferred to a phosphate buffered saline with 20% sucrose solution for at least three days prior to sectioning. 50-µm frozen sections were cut coronally or sagittally using a freezing microtome, and free-floating sections were processed for immunohistochemical labeling of GFP mounted on gelatin-coated glass slide and counterstained with a fluorescent blue counterstain. To visualize the injection site and axonal projections from viral transfected cells, sections were imaged at 10x using a Zeiss fluorescent microscope using Neurolucida software (MBF Biosciences, Williston, VT) to produce a whole brain reconstruction from tiled images of coronal or sagittal sections.

### Experimental design and statistical analysis

Data came from a total number of nine mice in the study. Eight mice were vgat-ires-cre, one mouse was C57BL/6J. Among the eight vgat-ires-cre mice used in the paper, five were male and three were females, the 1 C57BL/6J was male. We did not observe any systematic difference in behavioral performance between genders in this study. Four mice (vgat-ires-cre) were used for studying effects of 300-ms SC inhibition on detection performance; among them two were males, the other two were females. One mouse (C57BL/6J) was used for electrophysiological recordings. Eight (vgat-ires-cre) mice were used for studying effects of three 100-ms temporal epochs of SC inhibition on detection performance, and the same eight mice were used for studying effects of SC inhibition on psychometric curves. Five (vgat-ires-cre) mice were used for studying effects of orange light stimulation on detection performance, two were females and three were males. Five (vgat-ires-cre) mice were used for eye tracking while with SC inhibition, two were females and three were males.

For the psychometric data, results were pooled across sessions for each mouse to tabulate the overall lick probability for each amplitude of orientation change (including no change) separately for left and right visual fields and for with and without optogenetic stimulation. The lick probabilities were fitted with a cumulative Gaussian function using Psignifit3 (Wichmann and Hill, 2001). The fitted function had four parameters: mean, standard deviation of the Gaussian, bias (lower asymptote), and lapse rate (upper asymptote). Based on the standard deviation, we calculated the just-noticeable difference (JND), defined as the standard deviation multiplied by √2; the JND indicates the minimum change in signal strength that increases the detection rate by at least 50% of the available response range, once response bias and lapse rate have been taken into account.

For experiments using a single value of orientation change, we also pooled across sessions to tabulate lick probability, and tabulated hit and false alarm rates based on the definitions of trial outcomes described for the behavioral tasks. Performance was then characterized by measuring sensitivity (d’) and criterion using methods from signal detection theory (Macmillan and Creelman, 2005), as follows: d’ = Φ^-1^ (H) – Φ^-1^ (F), criterion = - (Φ^-1^ (H) + Φ^-1^ (F)/2, where Φ^-1^ is the inverse of normal cumulative distribution function, H is the hit rate and F is the false alarm rate. The 95% confidence intervals of hit and false alarm rates were generated with the binofit function in Matlab, which uses the Clopper-Pearson method to calculate confidence intervals. The 95% confidence intervals of d’ and criterion were generated with bootstrapped resampling.

Statistical analyses were conducted in Matlab using the statistics and machine learning toolbox and Prism 7 (GraphPad), and statistical significance was accepted as p<0.05. Paired sample nonparametric Wilcoxon signed-rank tests were performed to compare the cross-subject effect of SC inhibition on hit rate, false alarm rate, reaction times, d’ and criterion across the population of mice, unless stated otherwise. *Chi-square* tests were performed to compare the within subject effect of SC inhibition on hit rate. Nonparametric Wilcoxon rank-sum tests were performed to compare the cross-subject effect of 300-ms SC inhibition on hit rate and reaction times, and within-subject effect of SC inhibition on reaction times. Two-way ANOVAs were used to assess the interaction between SC inhibition during epoch 2 and spatial location for reaction times, using (inhibition vs no inhibition) and (contralateral vs ipsilateral) as two independent factors, the interactions were further tested with *post hoc* multiple comparisons using Bonferroni methods. Paired sample nonparametric Wilcoxon signed-rank tests were performed to compare the across-subject effect of SC inhibition on fitting parameters of psychometric curves (threshold, JND, lapse rate, guess rate). Nonparametric Wilcoxon rank-sum tests were performed to compare the cross-subject effect of stimulus competition on the magnitude of SC inhibition caused changes in fitting parameters of psychometric curves (threshold, JND, lapse rate, guess rate). The value of n reported in the figures and results indicates the number of animals. Error bars in figures indicate 95% confidence interval of the median or mean, unless indicated otherwise.

## Code Accessibility

All of the data were acquired and initially processed using custom scripts written in Matlab (The Mathworks, Natick, MA, USA). The Matlab code and data that support the findings of this study will be made available from the corresponding author upon reasonable request.

## Results

### SC inhibition caused deficits in change detection for contralateral hemifield

We inhibited neuronal activity in the SC by targeted optogenetic activation of GABAergic inhibitory neurons during a visual change-detection task (Figure 1A); we will refer to this optogenetic manipulation as “SC inhibition” throughout the study. Head-fixed mice were trained to detect near-threshold changes in the orientation of vertical Gabor patches centered in the left and right visual displays, using a task described previously (Wang et al., 2018; Wang and Krauzlis, 2018). Each trial was self-initiated, and mice progressed through a sequence of epochs based on a randomized distance traveled on the wheel (Figure 1C).

**Figure 1.**
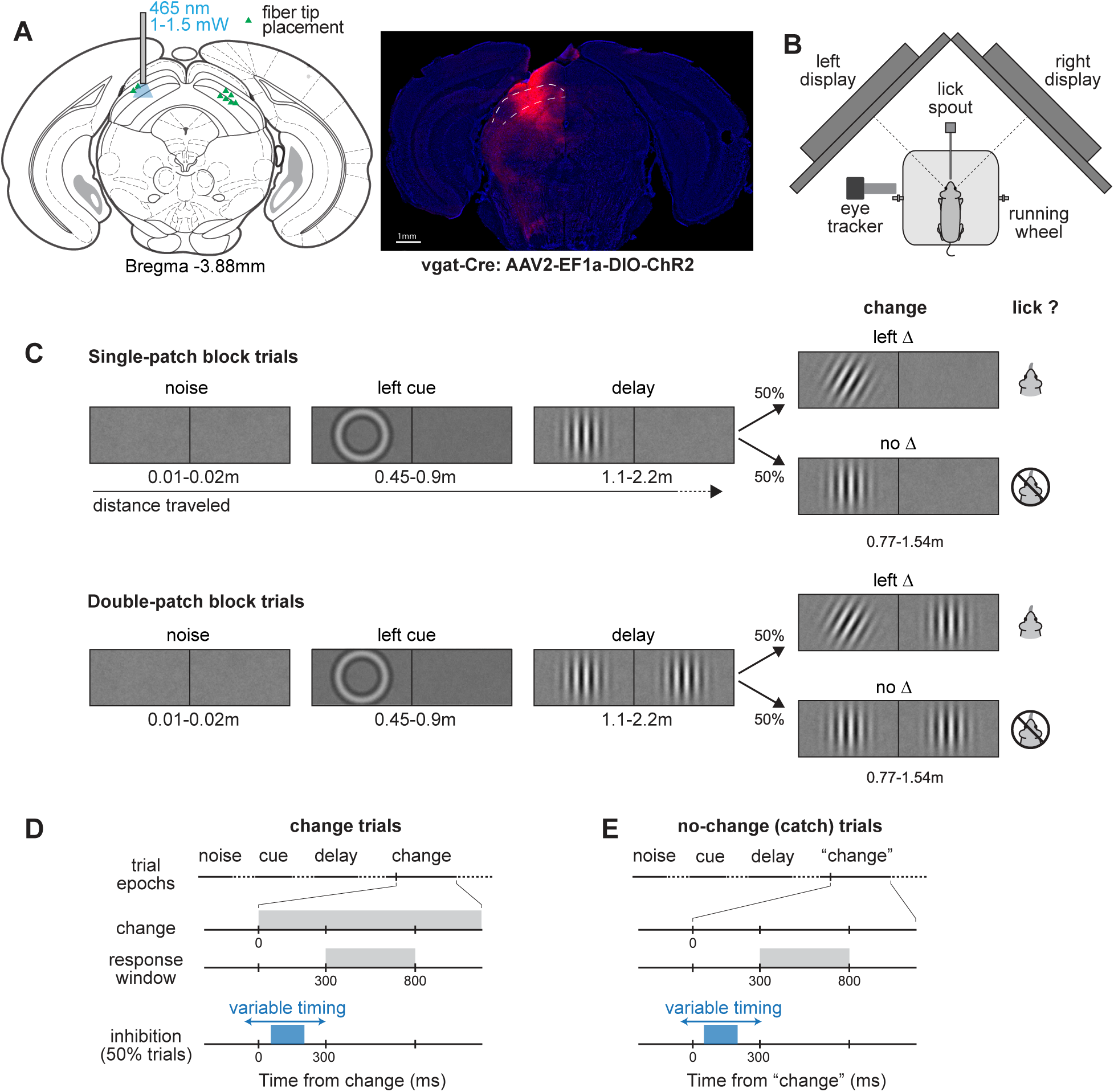
Schematics of orientation change detection task and superior colliculus (SC) optogenetics stimulation in mice. A) Localization of optogenetic manipulation in the SC. Left: stereotactic placement of optical fiber for optical stimulation. Right: coronal section from a sample vgat-cre mouse showing viral expression in the SC. B) Top-down view of the behavioral apparatus showing the placement of the lick-spout, and a head-fixed mouse on a Styrofoam wheel viewing two visual displays. C) Sequence of visual epochs during the detect task for two trial types, illustrating a single-patch (upper panel) and a double-patch (lower panel) trial with left-side orientation change. The length of each epoch was defined as distance mouse traveled through the trial. D and E) Timeline of events and response windows in change (D) and no-change (E) trials. Trial outcomes were defined based on a 500-ms response window for licks, which started 300ms after the orientation change; extraneous licks triggered trials to abort. The optogenetic stimulation window varied between across experimental conditions, but all ended before the beginning of response window.

The causal role of SC in detection performance was investigated by unilateral SC inhibition during various temporal intervals before the response window (Figure 1D). We first assessed whether the SC in mice was involved in the orientation-change detection by applying SC inhibition during a 300-ms interval spanning the time from the visual change to the onset of the response window (Figure 2A). Without SC inhibition, a cohort of 4 mice achieved high levels of performance for detecting 15° deg orientation changes (average hit rate of 90.3% and average false alarm rate of 10.2%). SC inhibition during the 300-ms interval significantly reduced hit rate specifically for orientation changes in the visual hemifield contralateral to the site of SC inhibition. Reductions were evident in both single-patch trials (Figure 2B), where contralateral hit rates with inhibition were all substantially below unity line (no inhibition: 86.9 ± 6.6%, mean ± 95%CI; with inhibition: 64.7 ± 13.0%, p = 0.057, Wilcoxon rank-sum test for the population; 4 out of 4 individual mice had significant reductions in hit rates with SC inhibition by Chi-square test, p<0.05) and double-patch trials (Figure 2C, no stimulation: 91.7 ± 4.1%, with stimulation: 60.2 ± 12.2%, p = 0.029; 4 out of 4 mice significant by Chi-square test, p<0.0001). In contrast to these strong contralateral effects, detection of orientation changes in the ipsilateral visual hemifield was unchanged by unilateral SC inhibition for both single-patch trials (Figure 2B, p = 0.57, Wilcoxon rank-sum test) and double-patch trials (Figure 2C, p = 0.99), consistent with the retinotopic organization of the SC. The false alarm rates were not markedly changed by the SC inhibition for either the contralateral or ipsilateral trials, as evident in Figure 2B-C (contralateral single-patch: p = 0.99, Wilcoxon rank-sum test; double-patch: p = 0.31; ipsilateral single-patch: p = 0.89, double-patch: p = 0.89). SC inhibition also significantly increased reaction times for detecting contralateral orientation changes, as evident in the results from double-patch trials shown in Figure 2E (no inhibition 404.1 ± 32.4 ms, with inhibition 462.8 ± 39.3 ms, p = 0.11, Wilcoxon rank-sum test for the population; 4 out of 4 mice had significant increases in reaction time by SC inhibition with p<0.01 in rank-sum test), and less obviously for single-patch trials (Figure 2D, no inhibition: 419.1 ± 54.0 ms, with inhibition: 462.2 ± 40.9 ms, p = 0.31; 3 out of 4 mice had significant increases in reaction time with p<0.01 for rank-sum test). Reaction times for ipsilateral orientation changes were not altered by SC inhibition for either single-patch or double-patch trials, as shown in Figure 2D-E (single-patch: p = 0.89; double-patch: p = 0.89). These spatially specific deficits in response accuracy and reaction time demonstrate that SC inhibition during this 300-ms window disrupts choice behavior but does not identify which stage of processing (i.e., anticipation, visual processing, response preparation) was most strongly affected.

**Figure 2.**
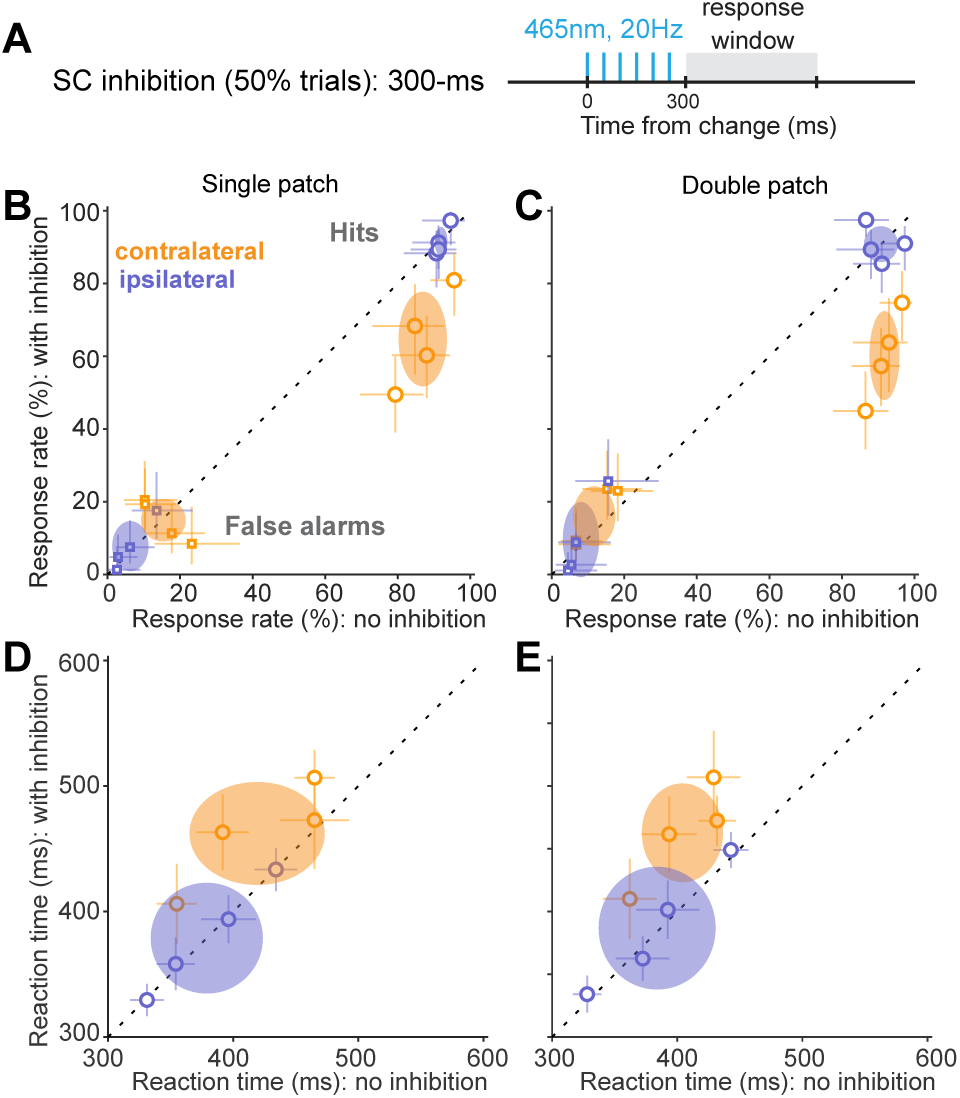
Unilateral SC inhibition impaired detection of contralateral orientation changes. A) Timing of the 300-ms optogenetic stimulation with respect to the orientation change. B) Single-patch trial hit rates (circles) and false alarm rates (squares) with versus without SC optogenetic stimulation for the 4 mice, plotted separately for contralateral (orange) and ipsilateral (blue) blocks. Open symbols are data from individual mice. Error bars indicate 95% CI. Ellipses are 95% CI of population mean. C) Same as B, but for double-patch trials. D) Reaction times with versus without SC optogenetic stimulation for single-patch trials. Open symbols are median reaction time from individual mice, error bars are 95% CI. Ellipses are 95% CI of population mean. E) Same as D, but for double-patch trials.

### Temporal precision of SC inhibition on detection performance

To identify which aspects of task performance were affected, we tested the effects of SC inhibition applied within more specific time windows. We defined three non-overlapping 100-ms temporal epochs based on the timing of SC neuronal responses to orientation changes (Figure 3A). “Epoch 1” (from −50 to 50 ms with respect to the orientation change) corresponded to the pre-change baseline of SC activity (Figure 3B); a behavioral effect caused by inhibition during this epoch might suggest the involvement of SC activity in anticipation during the task. “Epoch 2” (from 50 to 150 ms after the orientation change) overlapped with the phasic visual response of SC neurons to the orientation change (Figure 3E); a behavioral effect here would indicate the importance of SC visual processing for the perceptual decision. “Epoch 3” (from 150 to 250 ms after change onset) corresponded to the sustained activity after the initial visual burst (Figure 3H); a behavioral effect caused by SC inhibition during this epoch might indicate a role for the SC in response preparation or later stages of visual processing.

**Figure 3.**
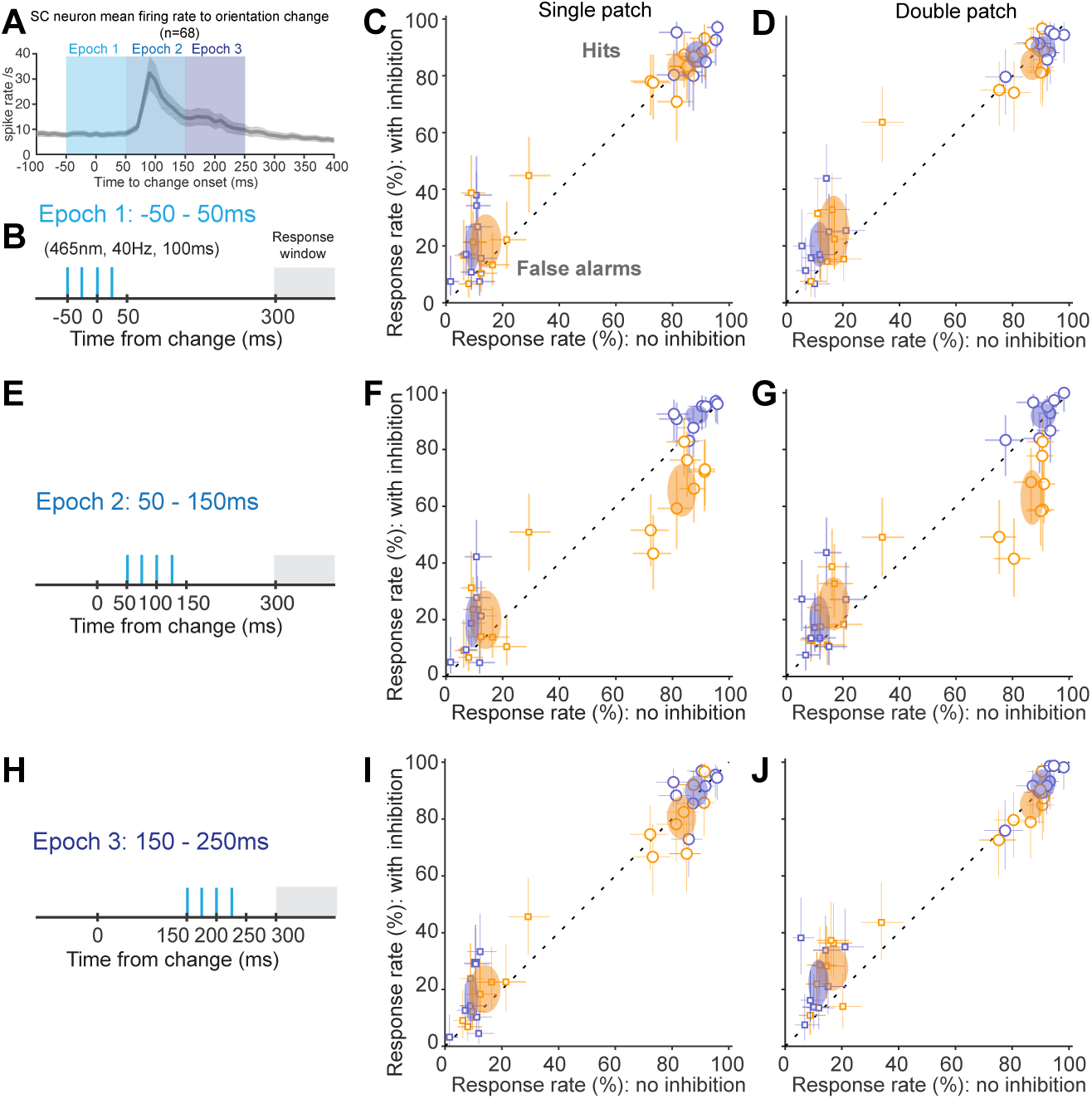
The effect of SC inhibition on detection performance had remarkable temporal specificity. A) Population PSTH of SC neuronal (n = 68, 28 from superficial layers, 40 from deeper layers) activity during presentation of a 30° orientation change in the contralateral visual field, aligned on the onset of the orientation change. Three 100-ms temporal epochs (shades of blue) indicate brief periods of SC inhibition used in the experiment. B) For epoch 1, 4 blue light pulses at 40 Hz were applied from −50 to 50 ms with respect to the orientation change. C) Single-patch trial hit rates (circles) and false alarm rates (squares) with versus without SC optogenetic stimulation applied during epoch 1 for 8 mice, plotted separately for contralateral (orange) and ipsilateral (blue) blocks. Open symbols are mean response rates from individual mice, error bars are 95% CI. Ellipses are 95% CI of population mean. D) Same as C, but for double-patch trials. E) For epoch 2, blue light pulses were applied from 50 to 150 ms with respect to the orientation change. F) Same as C, except that SC inhibition was applied during epoch 2. G) Same as F, but for double-patch trials. H) For epoch 3, blue light pulses were applied from 150 to 250 ms with respect to the orientation change. I) Same as C, except that SC inhibition was applied during epoch 3. J) Same as I, but for double-patch trials. Decreases in hit rate were only evident with SC inhibition when applied during epoch 2 for contralateral orientation changes (F, G).

SC inhibition during epoch 1 had little effect on task performance. Without SC inhibition, mice detected 12° orientation changes with average hit rates of 87.5% and false alarm rates of 12.8%. These rates were similar during epoch 1 inhibition for contralateral orientation changes, for both single-patch (Figure 3C, p = 0.74, Wilcoxon signed-rank test, comparing population hit rate with and without inhibition) and double-patch trials (Figure 3D, p = 0.15). Similarly, hit rates for ipsilateral orientation changes were also not altered by epoch 1 SC inhibition, as shown in Figure 3C-D (single-patch: p = 0.31; double-patch: p = 0.84). Epoch 1 inhibition had variable effects on false alarm rates. It did not significantly alter false alarm rates during contralateral blocks for either single-patch trials (Figure 3C, p = 0.25, Wilcoxon signed-rank test) or double-patch trials (Figure 3D, p = 0.11), but induced a small but significant increase of false alarm rates in both single-patch trials (Figure 3C, p = 0.035) and double-patch trials (Figure 3D, p = 0.016) during ipsilateral blocks. The absence of changes in hit rate by SC inhibition during this epoch suggests that SC activity during the delay period (as the mice might have been anticipating the orientation change) was not essential for task performance.

In contrast, inhibiting SC visual activity during epoch 2 caused profound deficits on detection performance. Epoch 2 inhibition significantly reduced hit rates for contralateral orientation changes in both single-patch (Figure 3F, p = 0.0078 Wilcoxon signed-rank test comparing between epoch 2 SC inhibition and no inhibition) and double-patch trials (Figure 3G, p = 0.0078). SC inhibition during epoch 2 did not have a significant effect on hit rates for ipsilateral orientation changes in either single-patch (Figure 3F, p = 0.055) or double-patch trials (Figure 3G, p = 0.55), consistent with the SC’s retinotopic organization. SC inhibition during this epoch also tended to increase false alarm rates for both contralateral (Figure 3F, single patch: p = 0.20, Wilcoxon signed-rank test; Figure 3G, double patch: p = 0.03) and ipsilateral (single patch: p = 0.045; double patch: p = 0.078) orientation changes. Consequently, sensitivity and response criterion also significantly changed with inhibition during this epoch. Sensitivity was significantly reduced with SC inhibition for detecting contralateral orientation changes (single patch, no inhibition: 2.12 ± 0.29, with inhibition: 1.31 ± 0.24, p = 0.0078, Wilcoxon signed-rank test; double patch, no inhibition: 2.13 ± 0.31, with inhibition: 1.01 ± 0.28, p = 0.0078) but not the ipsilateral ones (single patch, no inhibition: 2.61 ± 0.24, with inhibition: 2.41 ± 0.50, p = 0.38; double patch, no inhibition: 2.61 ± 0.31, with inhibition: 2.39 ± 0.52, p = 0.38). SC inhibition caused an increase in response criterion for contralateral changes (single patch, no inhibition: 0.06 ± 0.17, with inhibition: 0.25 ± 0.28, p = 0.11; double patch, no inhibition: −0.08 ± 0.10, with inhibition: 0.17 ± 0.24, p = 0.02) and a reduction for ipsilateral ones (single-patch, no inhibition: 0.06 ± 0.19, with inhibition: −0.25 ± 0.15, p = 0.02; double-patch, no inhibition: −0.09 ± 0.16, with inhibition: −0.31 ± 0.18, p = 0.07); the more lenient criterion was presumably an adaptation to the disruption of visual processing. These results demonstrate that the early visual responses of SC neurons to the orientation change are crucial for the ability of mice to perform this visual change-detection task.

SC inhibition during epoch 3 also had little effect on task performance. Hit rates were not significantly changed by SC inhibition during epoch 3 for either contralateral (Figure 3I-J, single patch: p = 0.38, Wilcoxon signed-rank test; double patch: p = 0.20) or ipsilateral orientation changes (single patch: p = 0.84; double patch: p = 0.47). Epoch 3 SC inhibition caused small but significant increases in false alarm rates for both contralateral (single-patch: p = 0.016; double-patch: p = 0.023) and ipsilateral (single-patch: p = 0.11; double-patch: p = 0.0078) blocks. The absence of effects on hit rates during this epoch suggests that this later stage of evoked SC activity was not essential for task performance and underscores the importance of the phasic visual response that immediately precedes this epoch.

SC inhibition also affected reaction times, but only for epoch 2. For epoch 1, there were no significant effects of SC inhibition on reaction times for either contralateral (Figure 4A, single-patch, p = 0.84, Wilcoxon signed-rank test; Figure 4B, double patch, p = 0.055) or ipsilateral orientation changes (Figure 4A, single patch, p = 0.38; Figure 4B, double patch, p = 0.11). Similarly, for epoch 3, there were no effects for contralateral (Figure 4E, single patch: p = 0.078; Figure 4F, double patch: p = 0.38) or ipsilateral changes (single patch: p = 0.055; double patch: p = 0.55). In contrast, SC inhibition during epoch 2 had a profound effect on reaction times. Reaction times for contralateral orientation changes with epoch 2 SC inhibition were significantly longer than without SC inhibition (Figure 4C, single patch: p = 0.016 Wilcoxon signed-rank test; Figure 4D, double patch: p = 0.0078). For ipsilateral orientation changes, the effects were more variable: SC inhibition did not alter reaction times for single-patch trials (Figure 4C, p = 0.99), but caused a small reduction for double-patch trials (Figure 4D, p = 0.039). Thus, epoch 2 SC inhibition during double-patch trials caused opposite effects on reaction time between contralateral and ipsilateral orientation changes (p = 0.038, two-way ANOVA), suggesting a cross-hemifield interaction in the presence of a visual distractor.

**Figure 4.**
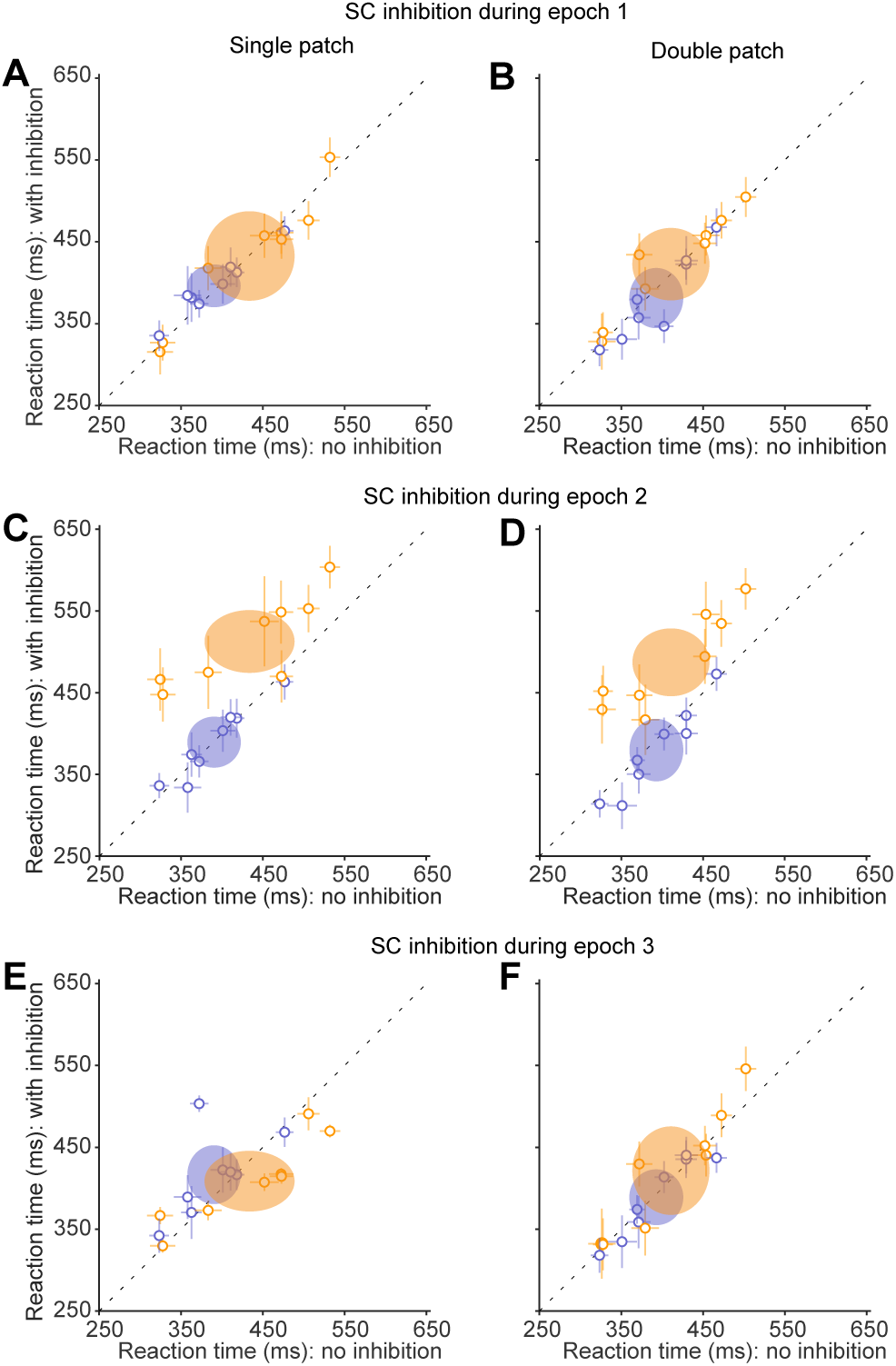
SC inhibition during the visual epoch increased reaction times for detecting contralateral orientation changes. A) Reaction times with versus without SC inhibition during epoch 1 for single-patch trials. Conventions are the same as in figure 2. B) Same as A, but for double-patch trials. C) Same as A, but with SC inhibition during epoch 2. D) Same as C, but for double-patch trials. E) Same as A, but with SC inhibition during epoch 3. F) Same as E, but for double-patch trials. Increases in reaction times were only evident with SC inhibition when applied during epoch 2 for contralateral orientation changes.

In summary, we found that the effects of inhibiting SC activity on hit rates and reaction times were specific to the 100-ms time epoch that matched visual-evoked phasic activity in the SC. These results indicate that SC visual responses are crucial for the ability of mice to detect the orientation changes in our task.

### Involvement of the mouse SC in visual selection revealed by psychometric curves

In the results presented so far, we showed that the ability to detect a 12° orientation change was markedly impaired when inhibition was applied during the initial SC visual response. There were also slightly larger effects on double-patch than single-patch trials, which might be expected given the widespread evidence in other species that the SC is involved in visual target selection (Krauzlis et al., 2013; Mysore and Knudsen, 2011). To test this further, we next investigated SC inhibition using a range of stimulus values and constructed psychometric curves. For these experiments, SC inhibition was applied only during epoch 2.

SC inhibition caused systematic changes in psychometric curves for contralateral orientation changes that were larger in the double-patch than single-patch condition. As illustrated by single-patch data from sample mouse #1 (Figure 5A), without SC inhibition (dark orange) the response rate on contralateral trials went from ∼10% to above 95% as the orientation change was increased from 4° to 20°. With SC inhibition (light orange), the curve was shifted rightward and/or downward, reaching an asymptote of only 75%. These effects of SC inhibition were more pronounced in double-patch trials (Figure 5B): the curve for SC inhibition was shallower and reached a lower asymptote of ∼65%. Thus, the detection deficits caused by SC inhibition appeared to be worse when a competing visual stimulus was present in the other hemifield. Similar effects were found across all the mice (Figure 5 C-D), albeit with some variability across animals, and an overall tendency for SC inhibition to increase false alarms. Specifically, the flattening of the psychometric curves caused by SC inhibition was more pronounced in the double-patch trials (Figure 5D, orange curves) than in the single-patch trials (Figure 5C).

**Figure 5.**
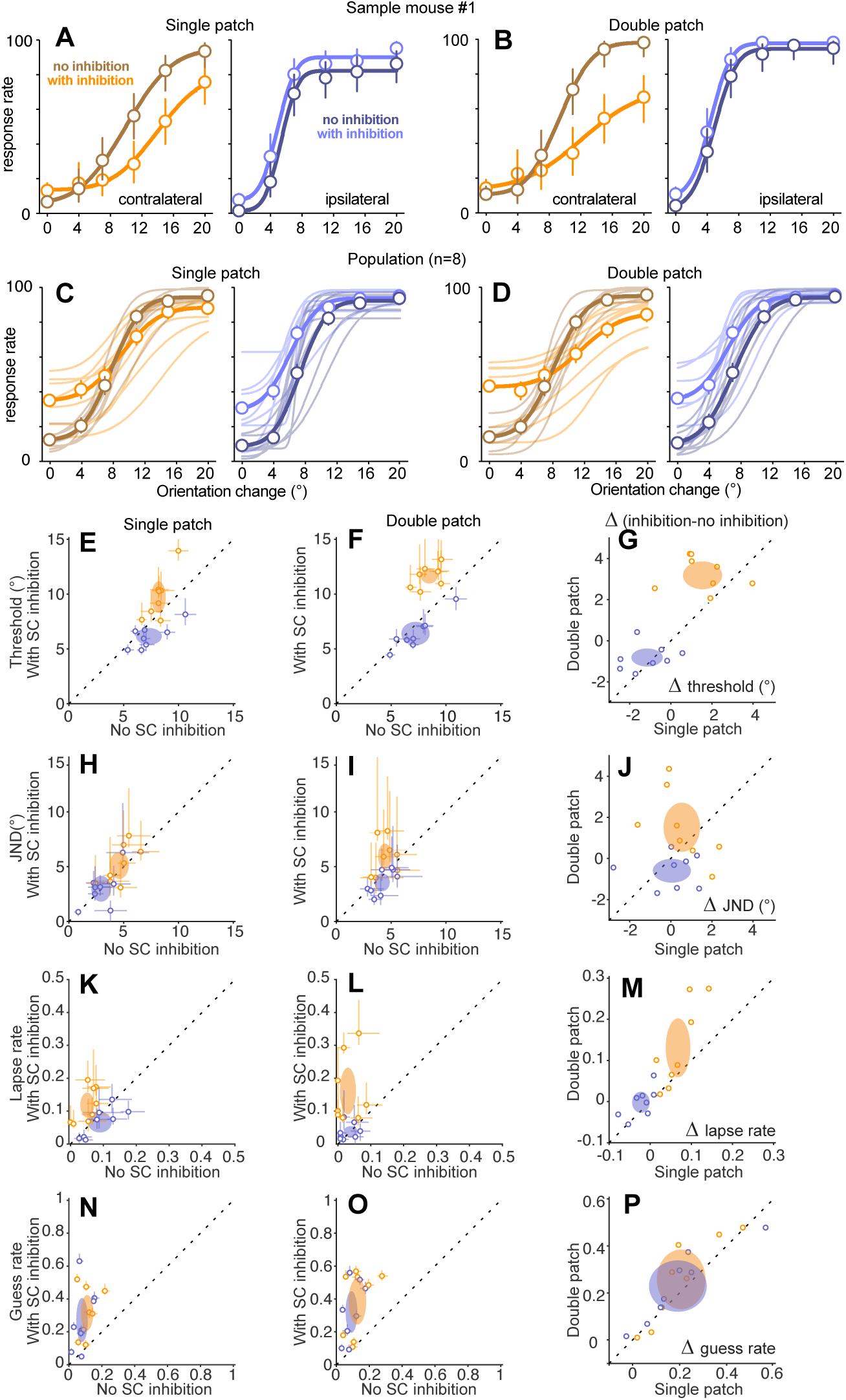
Effects of SC inhibition on psychometric curves. A-B) Psychometric data and curves from a sample mouse with and without SC inhibition applied during epoch 2 for single-patch (A) and double-patch (B) trials. Open symbols are mean response rates from individual mice, and error bars indicate 95% CI. Curves show fitted cumulative Gaussian functions. SC inhibition shifted psychometric curves downward for contralateral orientation changes. C-D) Psychometric curves across all mice (n = 8) with and without SC inhibition for contralateral (orange) and ipsilateral (blue) orientation changes. Thin lines are fitted curves from individual mice, and thick lines show population average, for single-patch (C) and double-patch (D) trials. E-F) Comparison of detection thresholds with versus without SC inhibition for contralateral (orange) and ipsilateral (blue) orientation changes, for single-patch (E) and double-patch (F) trials. Open symbols are median thresholds from individual mice, error bars are 95% CI. Ellipses are 95% CI of population mean. G) Comparison of the changes in thresholds (Δ threshold) induced by SC inhibition between single- and double-patch trials. H-J) Just-noticeable-differences (JND) with and without SC inhibition. K-M) Lapse rates with and without SC inhibition. N-P) Guess rates with and without SC inhibition. Other conventions as above.

To quantify how SC inhibition affected psychometric performance and identify whether those effects might depend on the presence of competing visual stimuli, we extracted four independent parameters of the cumulative Gaussian curves fitted to the data from individual mice, following methods described previously (Wang and Krauzlis, 2018). These four parameters were: 1) detection threshold, defined as the mean of the fitted Gaussian function, 2) just-noticeable-difference (or JND), the standard deviation of the fitted Gaussian function multiplied by √2, which is the minimum change in the visual orientation that would result in detection at least 50% of the time, 3) lapse rate, the upper asymptote of the cumulative function, and often attributed to non-sensory perceptual errors, and 4) guess rate, defined as the lower asymptote of the fitted function, and related to the subject’s response criterion or non-perceptual motor bias.

SC inhibition increased detection thresholds for contralateral orientation changes (Figure 5 E-F). These increases were significant for both single-patch (Figure 5E, p = 0.016, Wilcoxon signed-rank test) and double-patch trials (Figure 5F, p = 0.0078), and the effects on detection threshold were significantly larger for double-patch trials (Figure 5G, 3.25 ± 0.59°, mean ± 95%CI) than for single-patch trials (1.55 ± 0.95°, p = 0.039, Wilcoxon signed-rank test). Thus, SC inhibition decreased the likelihood of detecting smaller contralateral orientation changes and this effect was larger when a competing stimulus was present in the ipsilateral hemifield. For ipsilateral orientation changes, SC inhibition caused small decreases in detection thresholds during both single-patch (p = 0.039, Wilcoxon signed-rank test) and double-patch trials (p = 0.016) that did not differ in magnitude (Figure 5G, p = 0.55).

SC inhibition caused subtle changes in JND. For contralateral orientation changes, SC inhibition did not significantly alter JND during single-patch trials (Figure 5H, p = 0.25, Wilcoxon signed-rank test) but caused a small increase in JND during double-patch trials (Figure 5I, p = 0.042). This result suggests that suppressing the visual activity of SC neurons decreased the perceptual sensitivity for mice in our task, at least when a competing stimulus was present. However, the magnitude of contralateral JND increase caused by SC inhibition did not significantly differ between single-patch and double-patch trials (Figure 5J, p = 0.38). For ipsilateral orientation changes, SC inhibition did not significantly alter JND for either single-patch (Figure 5H, p = 0.46) or double-patch trials (Figure 5I, p = 0.11).

SC inhibition caused a spatially specific increase in lapse rates. For contralateral orientation changes, SC inhibition significantly increased lapse rates in both single-patch (Figure 5K, p = 0.0078, Wilcoxon signed-rank test) and double-patch trials (Figure 5L, p = 0.0078), and the size of the increase in lapse rates caused by SC inhibition in double-patch trials was greater than that in single-patch trials (Figure 5M, p = 0.039, Wilcoxon signed-rank test). Thus, the increased lapse rate caused by SC inhibition were amplified in the presence of a distractor in the ipsilateral hemifield, consistent with the idea that lapse rate is linked to uncertainty during the perceptual choice (Pisupati et al., 2019). These effects were also spatially specific, because lapse rates for detecting ipsilateral orientation changes were not significantly altered by SC inhibition for either single-patch (Figure 5K, p = 0.074) or double-patch (Figure 5L, p = 0.90).

Finally, SC inhibition caused nonspecific increases in guess rates. For single-patch trials, SC inhibition increased guess rates for detecting both contralateral (Figure 5N, p = 0.0078, Wilcoxon signed-rank test) and ipsilateral orientation changes (p = 0.016); the increases in guess rates for contralateral and ipsilateral changes were not significantly different (p = 0.88, two-way ANOVA with conditions of spatial locations and SC inhibition). Similarly, during double-patch trials, SC inhibition also increased guess rate for both contralateral (Figure 5O, p = 0.0078) and ipsilateral detections (p = 0.0078), also without spatial specificity (p = 0.78, two-way ANOVA). For both contralateral and ipsilateral trials, the increased guess rates caused by SC inhibition did not differ between single-patch and double-patch conditions (Figure 5P, contralateral: p = 0.15, Wilcoxon signed-rank test, ipsilateral: p = 0.11). These spatially non-specific increases of guess rate, along with the non-specific increase in false alarm rates when we inhibited SC activity during three temporal epochs (cf. Figure 3), are consistent with changes in response bias on trials with SC inhibition.

Inhibiting SC visual activity also significantly altered reaction times, but mostly for contralateral changes that were large enough to be detected (Figure 6). For contralateral orientation changes, SC inhibition increased reaction times for near-threshold and super-threshold orientation changes, but not for smaller orientation changes (Figure 6A-B). For ipsilateral orientation changes, the effect of SC inhibition on reaction times was smaller and more variable (Figure 6C-D).

**Figure 6.**
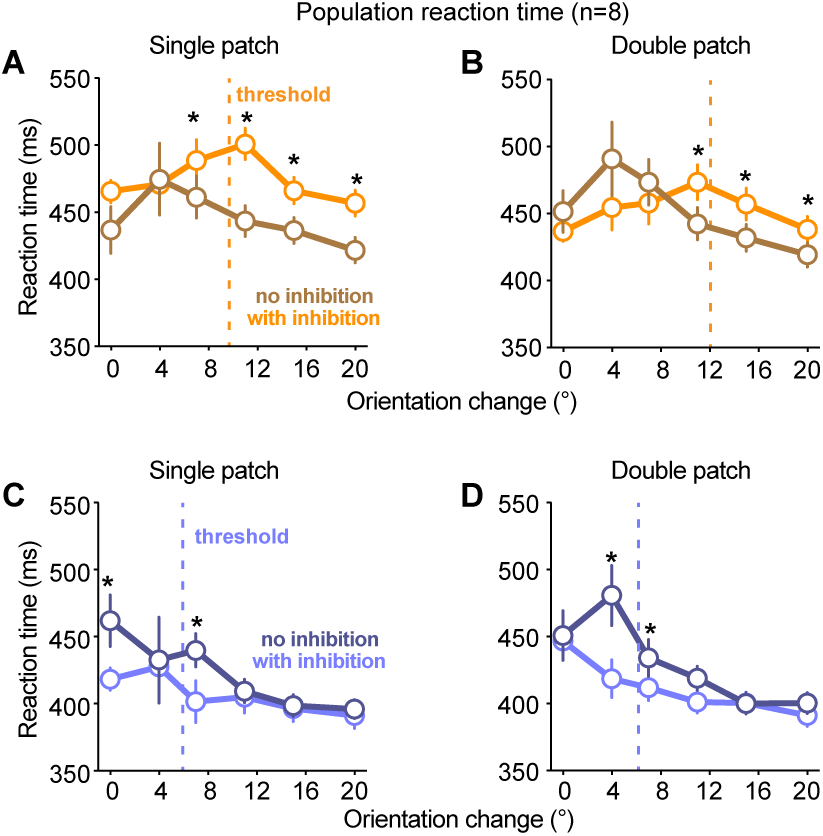
Effects of SC inhibition on reaction times for psychometric data. A-B) Population reaction times (n=8 mice) plotted as a function of orientation-change amplitude with (orange) and without (brown) SC inhibition for single-patch (A) and double-patch (B) trials. SC inhibition increased contralateral reaction times for orientation changes that were larger than the average threshold for detection (dashed line: population mean threshold with SC inhibition across mice). Symbols are median values and error bars indicate 95% CI. * indicates significant difference (p<0.05, Wilcoxon Signed rank test) in reaction time with and without SC inhibition at that particular orientation-change amplitude. C-D) Same as A-B, but for ipsilateral orientation changes. SC inhibition decreased ipsilateral reaction times for orientation-change amplitudes near threshold.

In summary, results from our psychometric analysis revealed that inhibiting SC visual activity for 100-ms caused a spatially selective deficit in several aspects of detection performance, including increases in detection thresholds and lapse rates for visual events represented by the affected SC neurons. The magnitude of these effects was larger in the presence of a competing visual distractor, consistent with a role of the SC in visual target selection as well as visual processing. In addition to deficits in the contralateral hemifield, inhibiting SC visual activity also caused a complementary improvement for some aspects of ipsilateral detection, including decreases in detection thresholds and decreases in reaction time, again consistent with a role for the SC in mediating competitive interactions across the visual field.

### Detection deficits not caused by retinal stimulation artifacts

One concern with optogenetic stimulation in mice is that scattered light from the stimulation might alter visual performance by activating the retina from inside the mouse’s head, rather than by changing the activity of neurons targeted for opsin expression. To assess whether the detection deficits we observed with optogenetic stimulation might be due to this type of nonspecific retinal activation, we performed a set of control experiments in a subset of vgat-cre mice used for data in Figures 2-6. For these controls (Figure 7A), we again stimulated the SC during epoch 2 using a 12° orientation change but instead of blue light (465 nm wavelength) we used orange light (620 nm wavelength). Orange light provides a strong test for possible retinal artifacts: the longer wavelength spreads further than blue light in brain tissue but does not activate ChR2 (Danskin et al., 2015).

**Figure 7.**
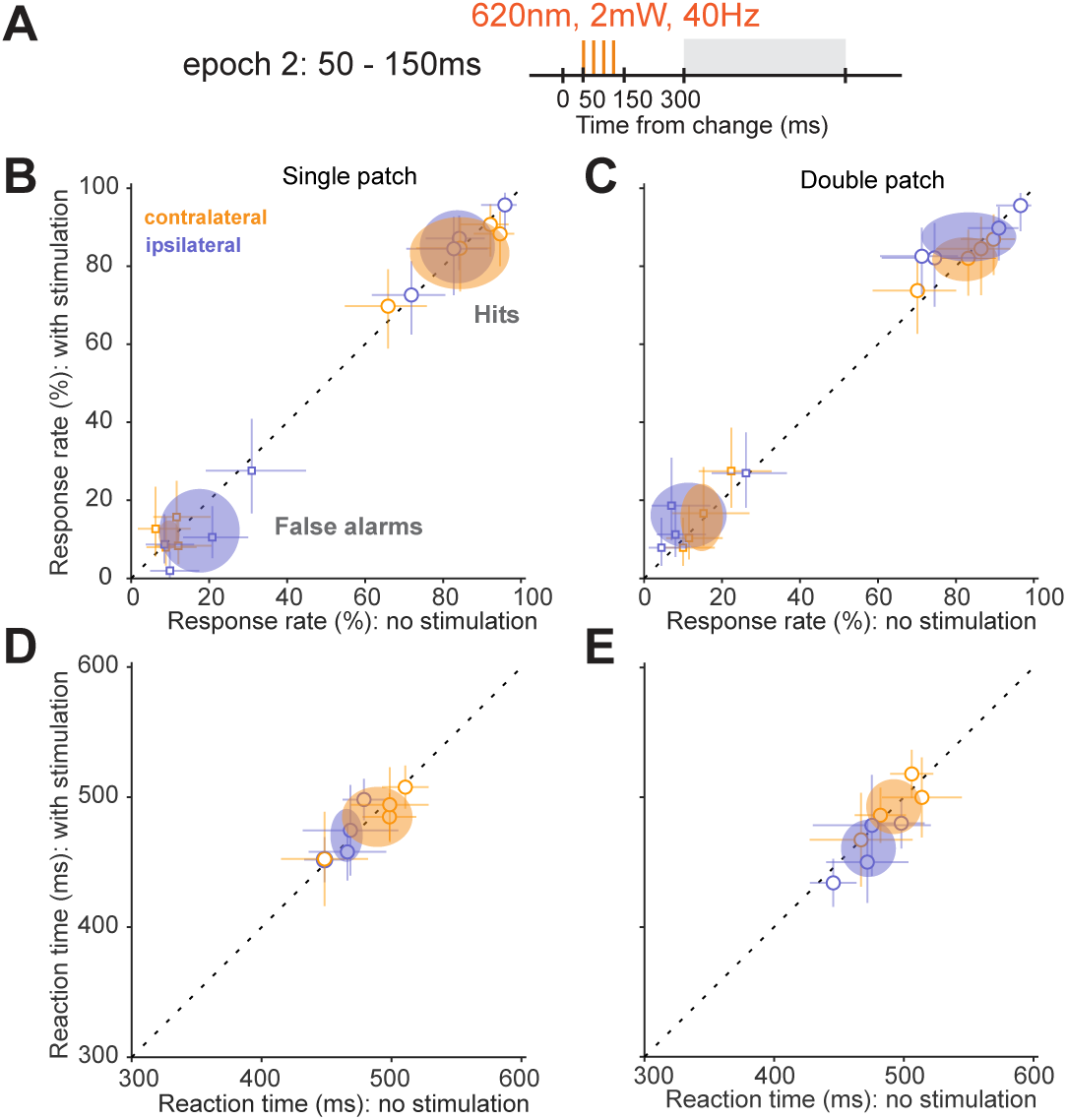
Orange light stimulation in the SC did not alter detection performance. A) Timing of the control 40-Hz orange-light (620 nm wavelength) stimulation applied in the SC during epoch 2, aligned to the orientation change. B) Single-patch trial hit rates (circles) and false alarm rates (squares) with versus without orange-light stimulation, plotted separately for contralateral (orange) and ipsilateral (blue) blocks. Conventions the same as in Figure 2. C) Same as B, but for double-patch trials. D) Reaction times with versus without orange-light stimulation for single-patch trials. Open symbols are median reaction time from individual mice, error bars are 95% CI. Ellipses are 95% CI of population mean. E) Same as D, but for double-patch trials.

Stimulation with orange light had no effect on task performance. For single-patch conditions, as shown in Figure 7B, orange-light stimulation caused no significant changes in hit rates compared to without stimulation for either contralateral (p = 0.88, Wilcoxon signed-rank test) or ipsilateral trials (p = 0.15). Similarly, false alarm rates were not altered by the orange-light stimulation (contralateral, p = 0.57; ipsilateral, p = 0.11). For double-patch conditions, as shown in Figure 7C, orange-light stimulation again did not change hit rate (contralateral, p = 0.72; ipsilateral, p = 0.28) or false alarm rate (contralateral, p = 0.65; ipsilateral, p = 0.14). Reaction times were also unaffected for both contralateral (Figure 7D, single-patch, p = 0.38; Figure 7E, double-patch, p = 0.88) and ipsilateral orientation changes (Figure 7D, single-patch, p = 0.38; Figure 7E, double-patch, p = 0.25). These results rule out the possibility that the deficits in task performance we observed during optogenetic stimulation were caused by artifactual stimulation of the retina.

### Detection deficits is not related to saccades

Another concern is that optogenetic inhibition in the SC might induce saccades (Wang et al., 2015), altering performance in the task by displacing the retinal image rather than by changing visual processing. To assess this possibility, we recorded the position of eye contralateral to the SC inhibition side during the visual epoch experiment (cf. Figures 3,4) in a subset of mice and measured the probability of saccades with and without SC inhibition. Since detection deficits were specific to SC inhibition during epoch 2, we focused on our analysis on that experimental condition.

Mice made very few saccades during our task, regardless of SC inhibition. As shown by a sample trial with saccades (Figure 8A), when saccades did occur, they were readily detected and tended to have larger horizontal than vertical components. Overall, the probability of saccades was below 3% throughout the trial, regardless of SC inhibition (Figure 8B), although SC inhibition did produce a slight increase in saccade probability (Figure 8B inset). To quantify this pattern, we compared saccade probability with and without epoch 2 inhibition during the time window from 50 to 800 ms with respect to the orientation change. In single-patch trials, SC inhibition did not change saccade probability on either change trials (p = 0.44, Wilcoxon signed-rank test, Figure 8C) or no-change trials (p = 0.31). In double-patch trials, SC inhibition caused a small but not significant increase in saccade probability for both change (p = 0.063) and no-change trials (p = 0.063). Thus, the SC inhibition generally did not have significant effects on saccade probability, and the change in probability of saccades was much smaller than the effect of SC inhibition on hit rates (cf. Figure 3). We conclude that the effects of SC inhibition on task performance was not an artifact of evoked saccades.

**Figure 8.**
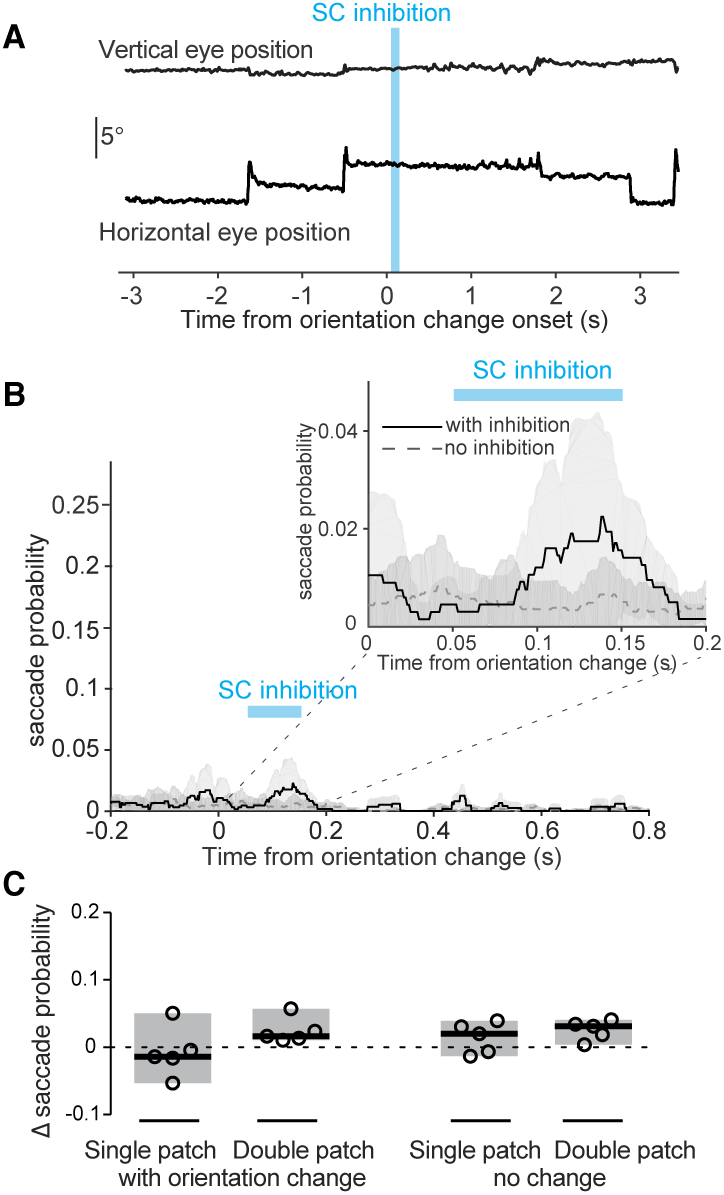
SC inhibition did not significantly alter saccade probability. A) Vertical and horizontal eye position traces from an example trial with SC inhibition applied during epoch 2 (blue bar). The orientation change occurred on the same side as the tracked eye. B) Saccade probability within each 1-ms time bin during double-patch trials with (solid black) and without (dashed gray) SC inhibition. The orientation change was again on the same side as the tracked eye and contralateral to the side with SC inhibition (n = 5 mice). Timing of the SC inhibition indicated by the blue bar. Inset shows an expanded view of saccade probability for the temporal interval between 0 and 200ms after the orientation change. C) Change in saccade probability (with SC inhibition minus no SC inhibition) during a 750-ms interval (from 50-ms to 800-ms after the onset of the change epoch). Showing 4 different trial types within contralateral cue blocks. Black lines indicate medians. Shaded regions indicate 95% CI.

## Discussion

Our findings highlight the importance of the SC in mouse visual perception. We found that the mouse SC is necessary for detection behavior during a visual orientation-change task. The most important temporal epoch was a 100-ms window that overlapped with the visual burst response of SC neurons. SC inhibition during this epoch caused profound deficits in detecting contralateral visual changes, including reduced perceptual sensitivity, increased detection threshold, prolonged reaction times and increased non-sensory lapse rates. In addition, deficits in detection were larger in the presence of a competing visual stimulus, consistent with a role for the SC in visual selection.

### Superior colliculus and early visual processing in mice

The retinocollicular pathway is the dominant visual pathway in mice. More than 95% of retinal ganglia cells (RGCs) in mice project to the superficial layers of the SC (Ellis et al., 2016; Hofbauer and Dräger, 1985), most RGCs that project to the lateral geniculate nucleus also project to the SC, and a significant proportion of RGCs project only to the SC (Ellis et al., 2016). In addition, visual features are represented in the mouse SC, including orientation (Wang et al., 2010), that are not present in the SC of primates but instead emerge in the visual cortex. Thus, the retinocollicular pathway in mice is responsible for much of the early visual processing that is associated with the retino-geniculate-cortical pathway in primates and other species.

Accordingly, although mouse V1 is important for visually guided behavior, it is not always essential and the effects of V1 lesions depend on the particulars of the visual task. Using a water maze task, complete removal of mouse V1 reduces visual acuity by about 50% for discriminating higher spatial frequency (>0.3cpd) gratings (Prusky and Douglas, 2004). However, the importance of V1 for visual detection and processing lower spatial frequencies is less clear. A recent study using low spatial frequency gratings similar to ours (∼0.1cpd) found that optogenetic inhibition of mouse V1 during a visual foraging task caused deficits in orientation discrimination performance (Resulaj et al., 2018) but left the detection of gratings intact. In contrast, another study using more sensitive psychometric measurement showed that inhibition of mouse V1 caused modest increases in detection thresholds for orientation or contrast changes (Glickfeld et al., 2013). Together these results indicate that mouse V1 is important for discriminating visual features, but less critical for visual detection especially at lower spatial frequencies.

In contrast, our results demonstrate that visual activity in the SC is crucial for detection performance in the mouse. The deficits caused by brief (100 ms) SC inhibition in our task were similar to, if not larger than, those caused by much longer V1 inhibition (seconds) spanning the full trial duration (Glickfeld et al., 2013). In fact, the modest detection deficit caused by mouse V1 inhibition might be mediated through the SC, as the gain of visual responses in SC can be modulated by inputs from V1(Zhao et al., 2014). Moreover, the effects of SC inhibition we observed exhibited remarkable temporal specificity. Detection performance was markedly impaired by inhibition applied during a temporal epoch that matched phasic visual activity in the SC, but not in the epochs immediately before or after. We do not conclude that SC activity in these other epochs is not important, since it is possible that deficits might emerge with stronger inhibition, but our results demonstrate the particular importance of visual activity in the SC for detection performance.

Visual information processed in the mouse SC is also passed along to visual cortical areas other than V1 during detection tasks. Outputs from mouse SC visual neurons modulate visual response properties of neurons in secondary visual cortex (Tohmi et al., 2014) through the lateral posterior nucleus of thalamus (Gale and Murphy, 2018; Tohmi et al., 2014; Zhou et al., 2018). The ascending projections from mouse SC appear to carry visual information distinct from V1 projections, particularly for visual motion. SC inhibition in mice has large effects on visual motion responses in postrhinal cortex (POR) (Beltramo and Scanziani, 2018), a higher order visual cortical area crucial for spatial navigation (LaChance et al, 2019), whereas inhibiting V1 has no effect on the motion response of POR neurons. In addition, SC lesions in mice also shift velocity tuning curves towards slower speeds in many secondary visual cortical areas (Tohmi et al., 2014). Thus, it is possible that the detection deficits caused by SC inhibition in our task, which used drifting visual gratings, could be due to disruptions in the processing of visual motion signals.

Alternatively, the perceptual deficits caused by SC inhibition could be due to a more generalized deficit in visual detection. Studies in primates have shown that SC inhibition causes deficits in detecting changes of visual features that are not represented by SC neurons (Herman et al., 2018). However, since neurons in the mouse SC and V1 are tuned to very similar visual features (Ito et al, 2017; Wang et al., 2010), to test whether the mouse SC serves as a general event detector would require additional experiments designed to test visual features that are not represented in the mouse SC, such as object identity.

### Superior colliculus and visual selection

We found SC inhibition in mice caused larger deficits in detection when a competing visual stimulus was present in the other visual hemifield, consistent with a role for the SC in mediating visual competition and selective attention (Krauzlis et al., 2013; Mysore and Knudsen, 2011). For example, perceptual deficits caused by reversible inactivation of the primate SC are small when only a single visual stimulus is placed in the affected part of the visual field, but much larger when competing distractors are included elsewhere in the visual field (Lovejoy and Krauzlis, 2009; 2017). In contrast, although we found larger deficits with competing stimuli (our double-patch condition), we also found substantial deficits even with a single stimulus placed in the affected visual field.

One explanation for this difference between our results and previous findings in primate SC is the relative importance of the SC to the rest of the visual system. In monkeys, the retino-geniculate-cortical route is the dominant visual pathway; very little early visual processing is done within the SC, where superficial layers receive retinal inputs only from less than 10% of RGCs (Perry and Cowey, 1984), and most perceptual and saccade related functions of primate SC involve its prominent intermediate and deep layers, where SC visual activity signal behavioral priority within a retinotopic spatial map (Basso and May, 2017; Boehnke and Munoz, 2008). In contrast, as described above, the mouse SC is the primary recipient of visual signals from the retina. Even though we mainly targeted GABAergic neurons in the SC intermediate layers, the strong GABAergic collateral projections from intermediate to superficial layers (Phongphanphanee et al., 2011) means that our optogenetic manipulation probably inhibited both superficial and deeper layers. The strong deficits in early visual processing caused by this inhibition probably limited our ability to measure effects on visual target selection. Given the relatively small size of mouse SC and the complexity of intracollicular circuit organization, isolating effects on visual selection might require targeting either inhibitory inputs to the intermediate layers or subsets of intermediate layer GABAergic interneurons that lack axonal collaterals to the superficial layers.

Selecting relevant visual targets among competing distractors is an important aspect of visual selective attention. We previously demonstrated that mice can use spatial cues to improve their visual perceptual sensitivity and adjust their detection thresholds in tasks similar to those used in primates (Wang and Krauzlis, 2018). The involvement of the mouse SC in visual selection revealed in the current study suggests that the SC could be an important brain structure for controlling visual selective attention in mice. The importance of the mouse SC in early visual processing also suggests that the changes in perceptual sensitivity during visual selective attention tasks (Wang and Krauzlis, 2018) could be due to modulation of the visual representations in SC superficial layers.

### Conclusion

Our current study reveals that the SC in mice plays an important causal role in visual perceptual decision-making, in addition to its established role in mediating innate responses to visual stimuli. The involvement of SC in our visual detection task was specific to a 100-millisecond interval coincident with the phasic visual response of SC neurons. In addition to its role in visual detection, the larger deficits found in the presence of a competing visual stimulus are consistent with a role for the mouse SC in visual target selection, although this effect was less prominent than that found in previous primate studies. Thus, our study highlights the importance of the SC in mouse visual perception and opens an avenue for using the mouse SC as a model system to study neural mechanisms of voluntary visual choices.

## Acknowledgements

the authors thank N. Nichols and D. Yochelson for technical support. The work is supported by the National Eye Institute Intramural Research Program at the National Institutes of Health.

